# Bacterial risk evaluation of widely consumed fast foods by the younger population in Bangladesh and its potential health effects

**DOI:** 10.1101/2023.07.30.551147

**Authors:** Lamia Bintay Masud, Md. Imran Hossain, Md. Omor Faruk, Md. Akber Subahan Mahbub Tuha, Sajidur Rahman Akash, Khondakar Raisul Islam, Md. Sahabuddin, Md. Sarafat Ali

## Abstract

Food-borne illness is a significant public health concern in Bangladesh, especially among the younger generation. They often consume spicy street foods, such as Fusca, Hog plum bedfellow, and Chotpoti. These foods can be contaminated with harmful bacteria, which can cause various diseases, including diarrhea, cholera, and dysentery. This study investigated the microbiological quality of three commonly consumed street foods in Bangladesh. We collected samples of Fusca, Hog plum bedfellow, and Chotpoti from different vendors in Dhaka and analyzed them in a laboratory. We found that all three of the food samples were contaminated with a variety of bacteria, including Bacillus spp., Haemophilus spp., Salmonella spp., Klebsiella spp., Staphylococcus spp., and Streptococcus spp. Some of these bacteria, such as Haemophilus spp., were resistant to the antibiotic ciprofloxacin. Our findings suggest that there is a high risk of food-borne illness associated with the consumption of street foods in Bangladesh. We urge vendors and consumers to take steps to improve food safety, such as proper food handling and cooking and clean water and utensils.

## Introduction

Street food refers to food and drinks that are prepared and sold by fast vendors or retailers, typically on the spot or in the same public place, for local consumption or later consumption without further preparation [1]. Vendors or manufacturers usually prepare street food in mobile food carts or food trucks, and it is intended to be consumed by customers. Street food is incredibly popular among consumers due to its delicious taste, affordability, nutritional value, and easy accessibility [2]. These affordable street foods serve as a significant source of nutrients for a large portion of the population [3]. Furthermore, selling street food in cities creates job opportunities and income for many people. Consumers highly value street food for its taste, affordability, and availability at all times [4], [5]. In Bangladesh, street food plays a crucial role in meeting the daily nutritional needs of citizens who have busy schedules and are unable to cook meals at home [6]. Therefore, due to its quick availability, street food becomes one of the primary dining options for city dwellers [7]. However, it is worth noting that street food sold during fast-paced environments often lacks proper sanitation, leading to high levels of contamination. In many cases, retail stores selling street food do not have access to running water, and hand washing is often done in a dish or bucket, sometimes without soap [8]. Insufficient food handling practices among vendors, who lack knowledge and skills in food safety management and formal food safety training, can result in contamination [9]. Numerous studies have reported that street food, especially in densely populated countries, can harbor a wide range of bacteria [8]. Therefore, from a health perspective, the marketing of street food remains controversial and street food may play a significant role in the transmission of risky and potentially fatal foodborne infections [9].

Food sold by fast vendors in developing countries is a major contributor to foodborne diseases (FBD) [10]. Over the years, the consumption of non-homegrown foods has been linked to various foodborne illnesses [11]. These illnesses not only pose a significant threat to public health but also have economic consequences. Foodborne diseases encompass a wide range of illnesses caused by contaminants, viruses, bacteria, and chemicals in food, making it a major global public health issue [12]. Contamination can occur during storage, transportation, display, handling, preparation, or sale of food in unsanitary environments, lacking potable water, proper sanitation facilities, and adequate waste removal [9]. Individuals who regularly consume street food are at a higher risk of food poisoning, including diarrhea, cholera, and typhoid [13]. The unsanitary conditions in which street food is prepared, stored, and transported raise concerns about its microbiological quality [14]. Numerous studies have reported the presence of pathogenic pathogens such as Escherichia coli, Salmonella spp., Staphylococcus aureus, Bacillus cereus, Clostridium perfringens, and Vibrio cholera in street food [15]. Staphylococcus aureus and Clostridium perfringens, which cause stomach pains and diarrhea, as well as Bacillus cereus, which causes vomiting and diarrhea, are commonly found in street food. Additionally, Salmonella is responsible for typhoid fever, food poisoning, and inflammation of the gastrointestinal tract [16]. A comprehensive analysis of 332 studies conducted in 36 countries showed that antibiotic-resistant foodborne bacteria have become a global health concern. More than 19% of human clinical specimens tested positive for antibiotic-resistant foodborne pathogens, and these bacteria exhibited high levels of resistance to the antibiotics used in the study. Furthermore, approximately 90% of foodborne pathogenic bacteria were found to form biofilms. The economic impact of foodborne illnesses in low- and middle-income countries amounts to approximately $ 110 billion annually, and the development of antibiotic resistance in foodborne pathogens significantly affects public health. Antibiotic use in the global food production system remains prevalent, while antimicrobial resistance continues to accelerate among humans, animals, plants, and the environment [16].

In the Indian subcontinent, particularly in Bangladesh, the younger generations have a fondness for street foods such as Fusca, hog plum bedfellow, and Chotpoti, which they often enjoy during leisure time. However, our research indicates that common street foods are not always safe due to inadequate hygiene practices during their preparation, cooking, and serving. These foods may harbor various pathogenic bacteria, posing potential life-threatening risks to humans. This study aims to identify the types of pathogenic bacteria present in fast food and determine the level of antibiotic resistance among these pathogens by testing them against different generations of antibiotics used for treating human diseases.

## Materials and methods

The entire study was conducted in two steps. The first step involved isolating and identifying bacteria from the food samples using various microbiological tests, biochemical tests, and gram-staining. In the second step, the isolated bacteria were evaluated for antibiotic sensitivity.

### Sample Collection

Three food samples were collected from different fast food vendors in Ghonapara, Nabinbag, and Launch Ghat in Gopalganj district, Bangladesh (**Figure 1**). The food samples included Hog plum bedfellow (Spondias mombin), Fusca, and Chotpoti. Approximately 300g of each food sample was collected using the vendors’ serving utensils, wrapped in a parcel, and placed into sterile plastic bags. These samples were analyzed within 24 hours of collection.

**Figure 1.**
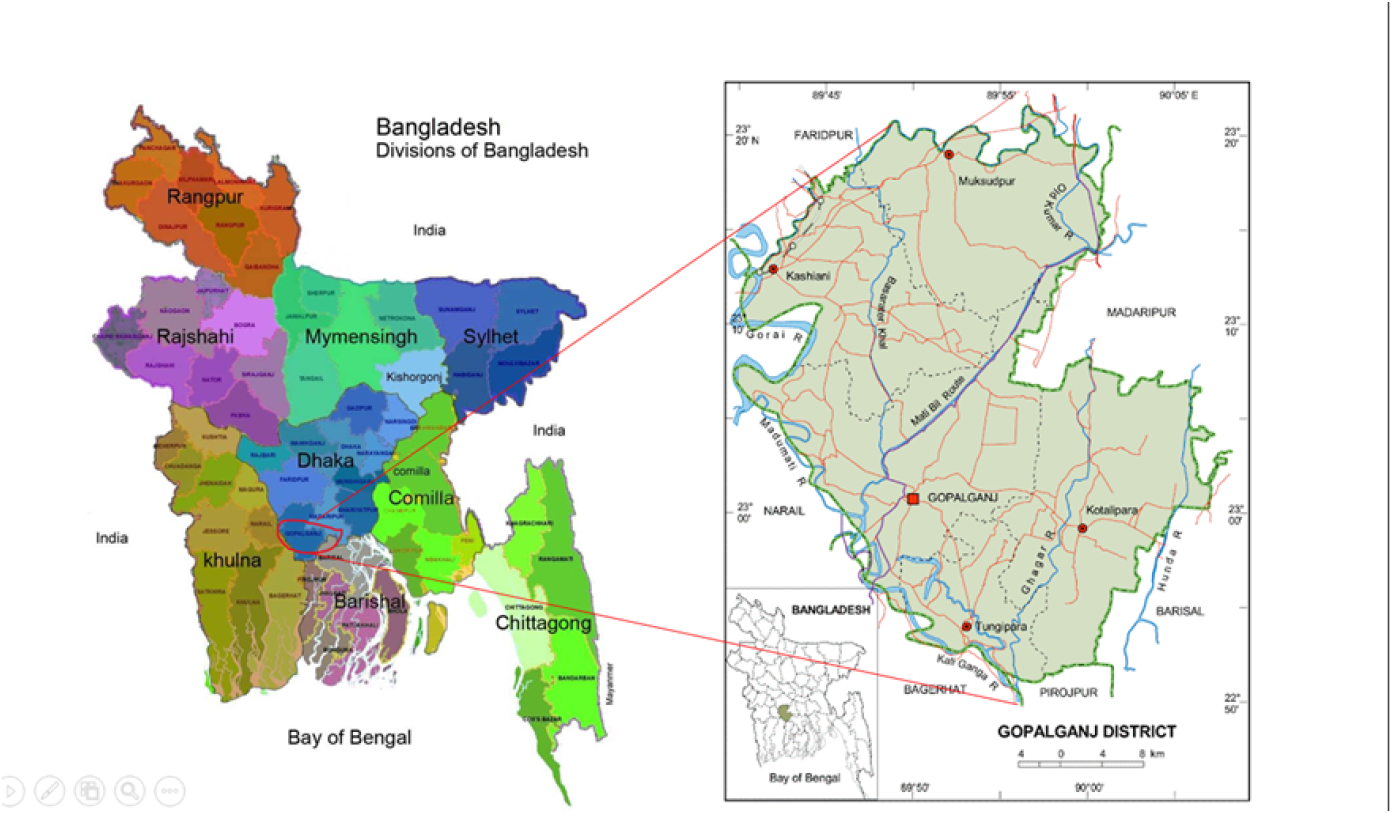
Map showing the location and outline of the study area.

### Sample Preparation

Approximately 15 grams of each street food sample (Hog plum bedfellow, Fusca, and Chotpoti) were uniformly homogenized in a mortar and pestle using a sterile diluent called phosphate-buffered saline (PBS). A homogenized suspension was created using the mortar and pestle. To create a standard solution, 1.0 gram of each homogenized sample was mixed with PBS. The solution was then serially diluted (1:10) to a dilution of 10-3 in test tubes, with 100 _μ_l of the stock solution added to 900 _μ_l of normal saline. Subsequently, 100 _μ_l of the homogenized samples from each dilution were spread on nutrient agar medium using a plastic spreader and incubated at 37°C for 24 hours. The diluted samples were spread as quickly as possible on the plate’s surface. After 24 hours of incubation, the plates were observed for bacterial growth.

A micropipette was used to transfer 0.1 ml of each tenfold dilution and spread it on a nutrient agar plate to calculate the total number of bacteria present. The diluted samples were spread across the plate’s surface as quickly as possible using a sterile glass spreader. This process was repeated for each dilution. The plates were then incubated at 37°C for 24 hours. The total bacterial count was determined as the number of colony-forming units (CFU) per gram of the tested sample.

### Bacteria Identification

The cultural examination of street food samples (Hog plum bedfellow, Fusca, and Chotpoti) for bacteriological study was conducted following the standard method provided by the International Commission on Microbiological Specifications for Foods [16]. Pure colonies were stained with a gram stain and examined under a microscope (UNITRON 14711-PS, USA) for morphological and staining characteristics. Various biochemical tests, such as indole, MIU, TSI, methyl red, Voges-Proskauer, citrate utilization, and urea utilization, were used for the identification and verification of these bacteria.

### Antibiotic Susceptibility Test

To determine the drug sensitivity and resistance patterns of the isolated bacterial strains, commercially available antibiotic discs (6 mm in diameter) were used, including Ciprofloxacin (5 μg/disc), Cefixime (5 μg/disc), Azithromycin (30 μg/disc), and Gentamicin (10 μg/disc). These antibiotics were selected based on their usage in the treatment of bacterial infections in patients from Gopalganj, Bangladesh. The Kirby-Bauer disc diffusion technique was employed to assess antibiotic resistance, following the guidelines of the Clinical and Laboratory Standards Institute [17]. After an overnight incubation at 37°C, the diameter (in millimeters) of the zones of inhibition around each antimicrobial disc was measured, recorded, and classified as resistant (R), intermediate (I), or sensitive (S).

## Results

### Isolated bacteria from samples

After 24 hr of incubation, we found there were numerous bacterial colonies in the nutrient agar plates (**Figure 2**). In the samples of Hog plum bedfellow, we found only five identical and a few confluent bacterial colonies; however, in Fusca, there were a significant number of confluent colonies as well as around 90 bacterial identical colonies were found. On the other hand, Chotpoti had a significant number of bacterial colonies, which is around 3-300 colonies per gram.

**Figure 2.**
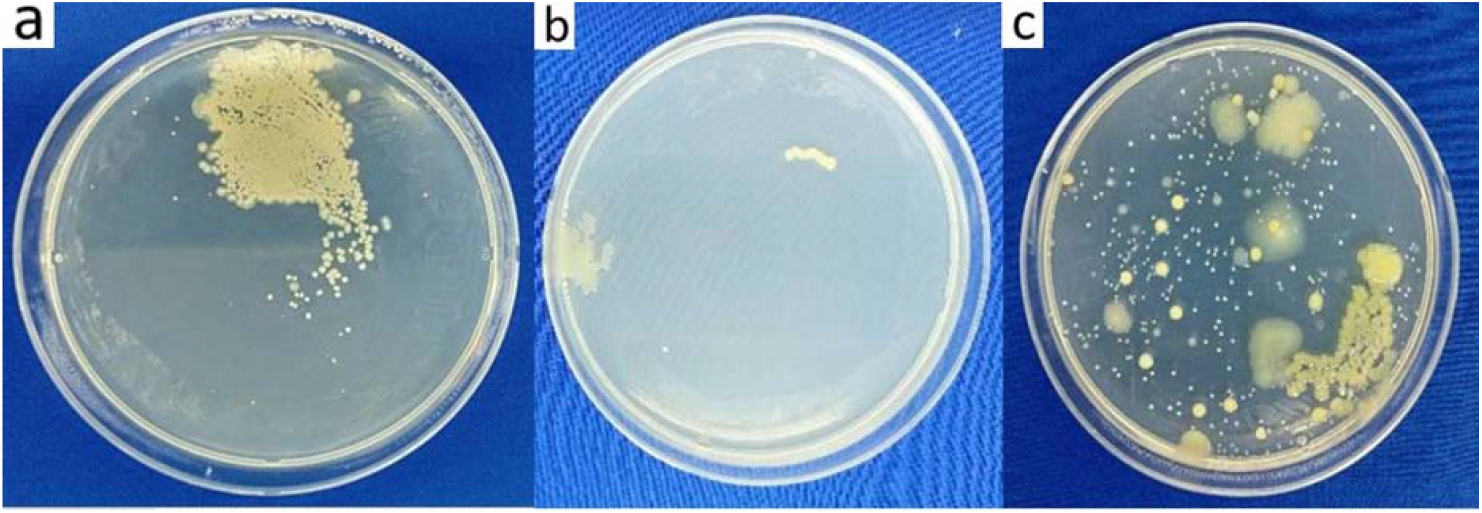
Isolation of bacterial colonies from three types of street food samples. a: Chotpoti; b: Hog plum, and c: Fusca.

**Figure 3.**
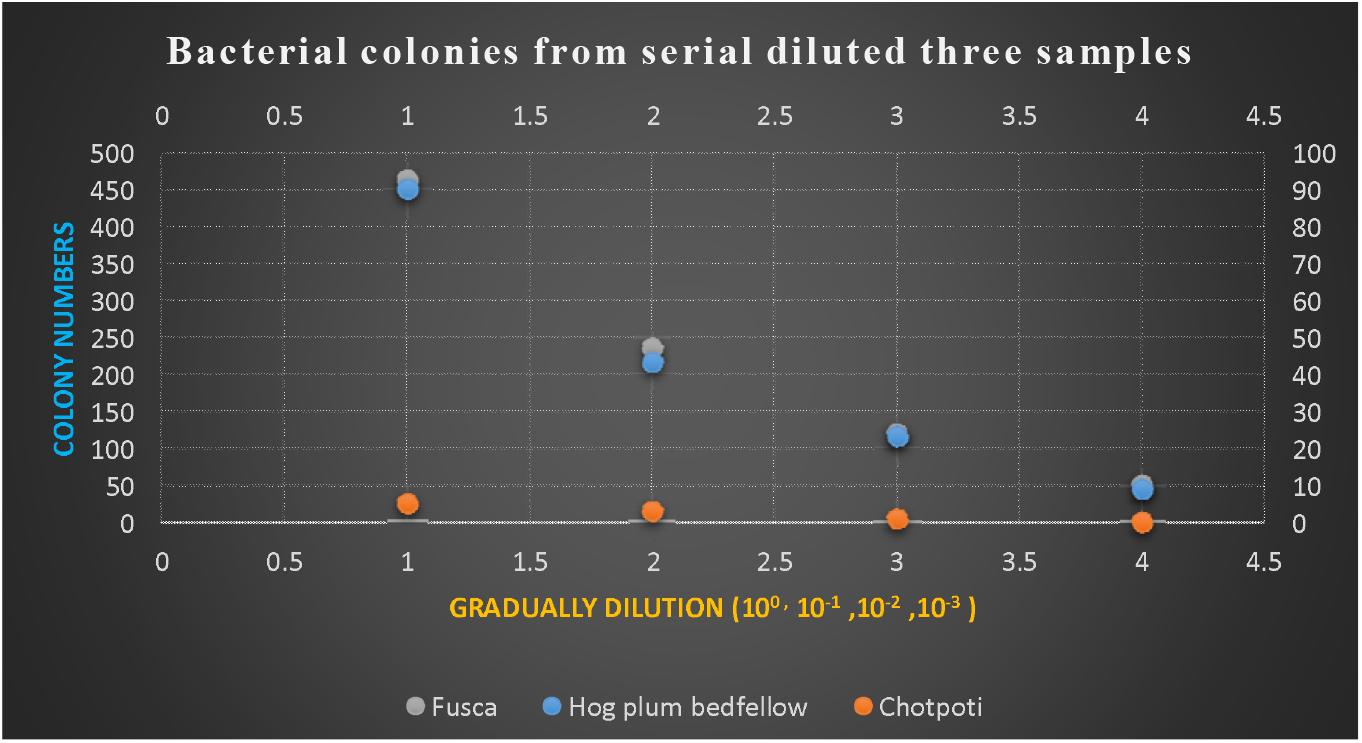
Scatter plot of bacterial colonies from serial diluted three samples (100, 10-1, 10-2 & 10-3).

### Total viable count

We have presented total viable cells in Table 1, where Fusca contain the highest number microbes nearly 37.3% and 45.8% than Chotpoti and Hog plum bedfellow. On the other hand, Hog plum bedfellow contained second position among these food items, around 8.5% higher Hog plum bedfellow.

**Table 1:**
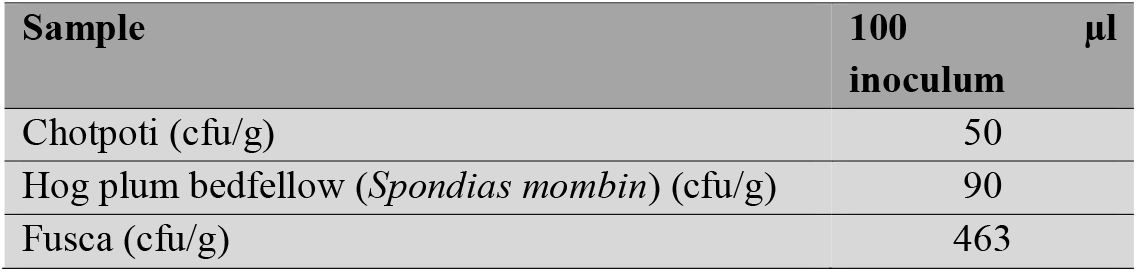
Total viable count of bacteria from collected three samples.

### Identification of associated bacteria

#### Characteristics by morphological study and gram staining

There were six types of bacterial species (Bacillus spp., Haemophillus spp., Salmonella spp., Klebsiella spp. Staphylococcus spp., Streptococcus spp.) identified on the basis of morphological study under the microscopic view along with Gram’s staining and different types of biochemical tests. In Chotpoti, there were two types of bacteria (**Figure 4**) (a) Bacillus spp., (b) Salmonella spp. Only one type bacteria Staphylococcus spp. was isolated from Hog plum bedfellow (**Figure 5**). On the other hand, we found four types of bacteria, such as (a) Klebsiella spp., (b) Haemophillus spp., (c) Streptococcus spp. and (d) Staphylococcus spp. from Fusca (**Figure 6**).

**Figure 4.**
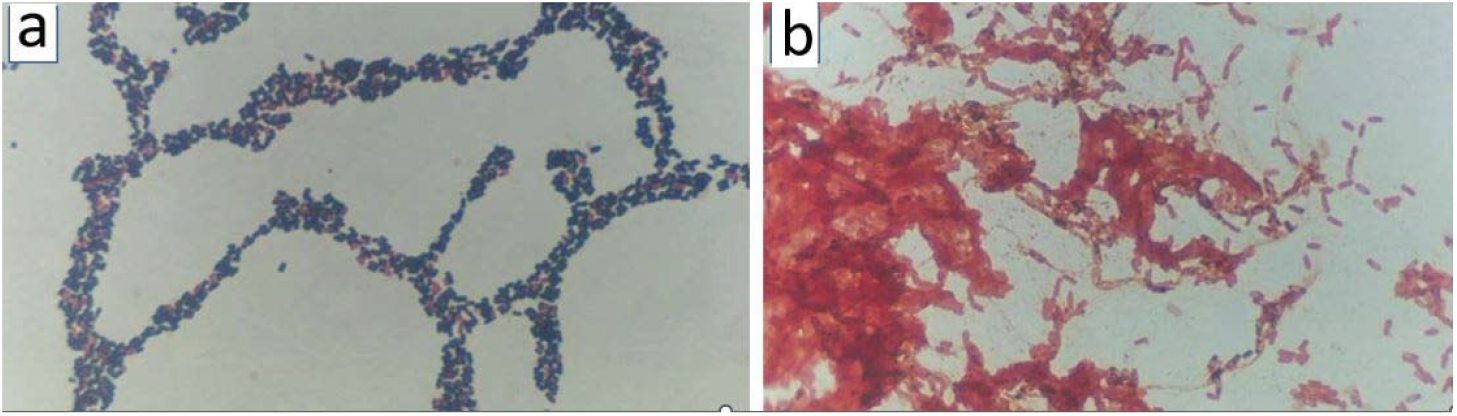
Light microscopy photograph at 100X magnification of isolated bacteria from Chotpoti (a) Bacillus spp., (b) Salmonella spp.

**Figure 5.**
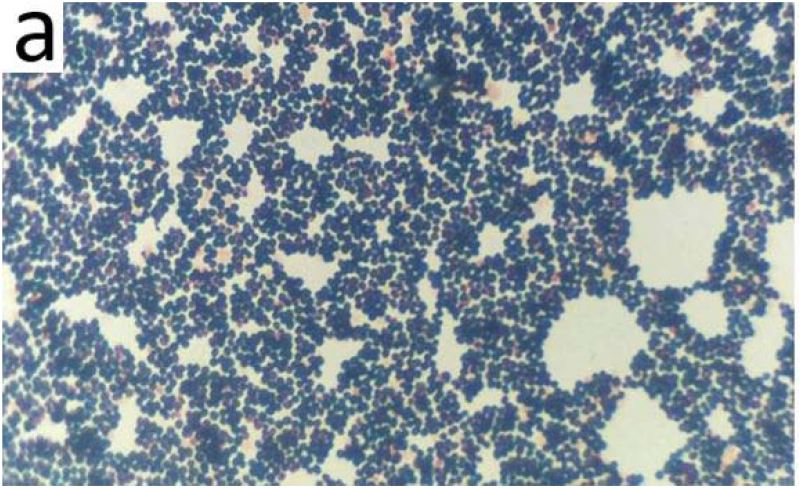
Light microscopy photograph at 100X magnification of isolated bacteria from Hog plum (a) Staphylococcus spp.

**Figure 6.**
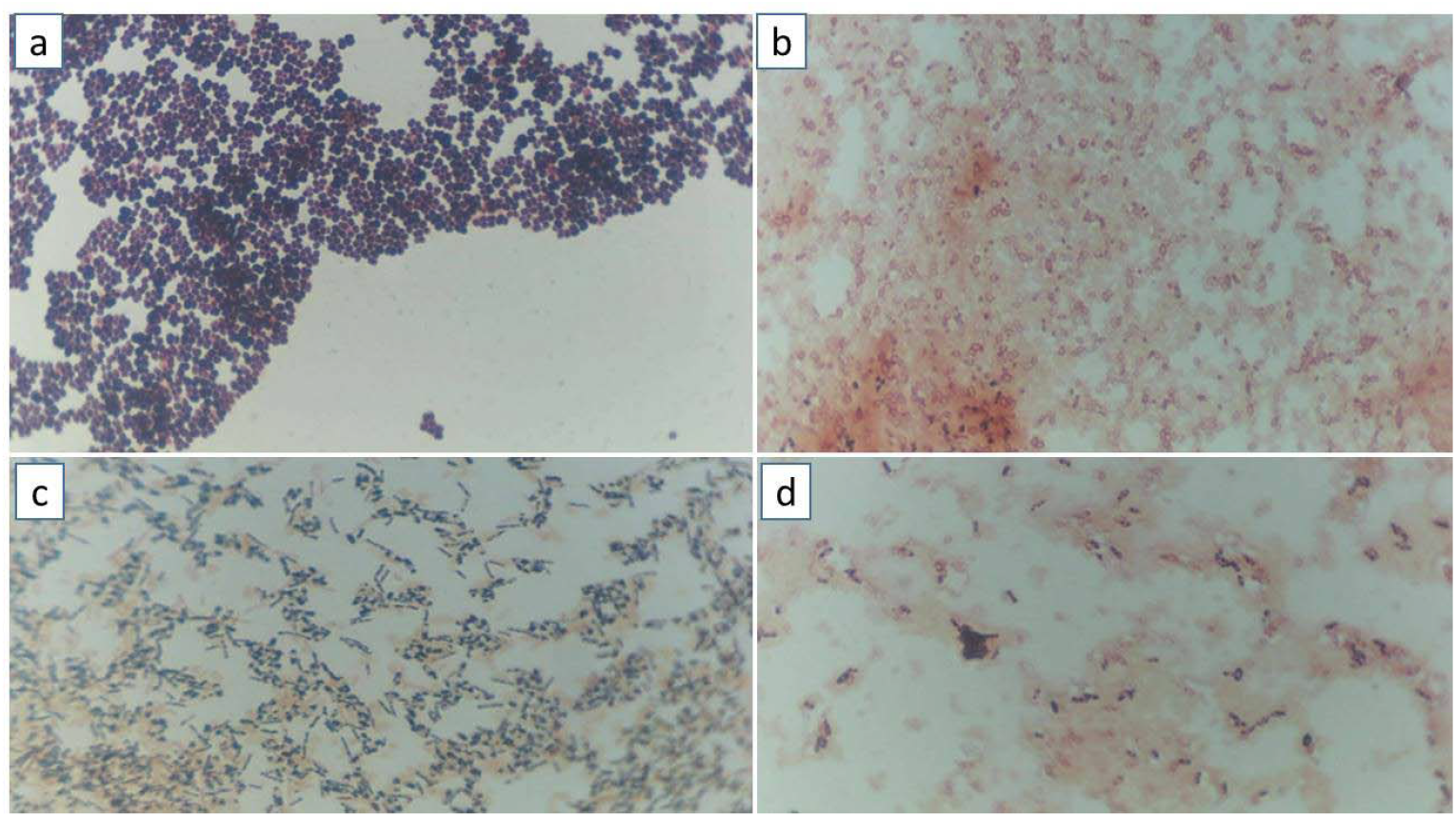
Light microscopy photograph at 100X magnification of isolated bacteria from Fusca; (a) Klebsiella spp., (b) Haemophillus spp., (c) Streptococcus spp. and (d) Staphylococcus spp.

The morphological and staining traits of bacteria observed in isolated samples under a microscope are presented in **Figures 4, 5, 6** and **Table 1**.

#### Characteristics by biochemical tests of isolated samples

The gram-staining test was performed for the microscopic characterization of bacteria and the results has been presented in the **Table 2**. The identified isolates were further confirmed by seven biochemical tests, such as indole, MIU, TSI, methyl red, Voges-Proskauer, citrate utilization, and urea utilization (**Table 3**).

**Table 2.** Microscopic characterization of isolated bacteria after gram staining.

| Sample | Microscopic characterization | Gram's reaction | Identified isolates |
| --- | --- | --- | --- |
| Fusca | Non-motile, encapsulated, lactose-fermenting, facultative anaerobic, rod-shaped bacterium | Negative | *Klebsiella* spp. |
|  | Nonmotile, Rod shaped | Negative | *Haemophilus* spp. |
|  | Coccoid, nonmotile and non-spore forming. | Positive | *Streptococcus* spp. |
|  | Nonmotile, non-spore forming, spherical shape | Positive | *Staphylococcus* spp. |
| Hog Plum | Nonmotile, non-spore forming, spherical shape | Positive | *Staphylococcus* spp. |
| Chotpoti | Rod-shaped, endospore-forming aerobic or facultatively anaerobic | Positive | *Bacillus* spp. |
|  | Straight rod shaped, motile | Negative | *Salmonella* spp. |

**Table 3.**
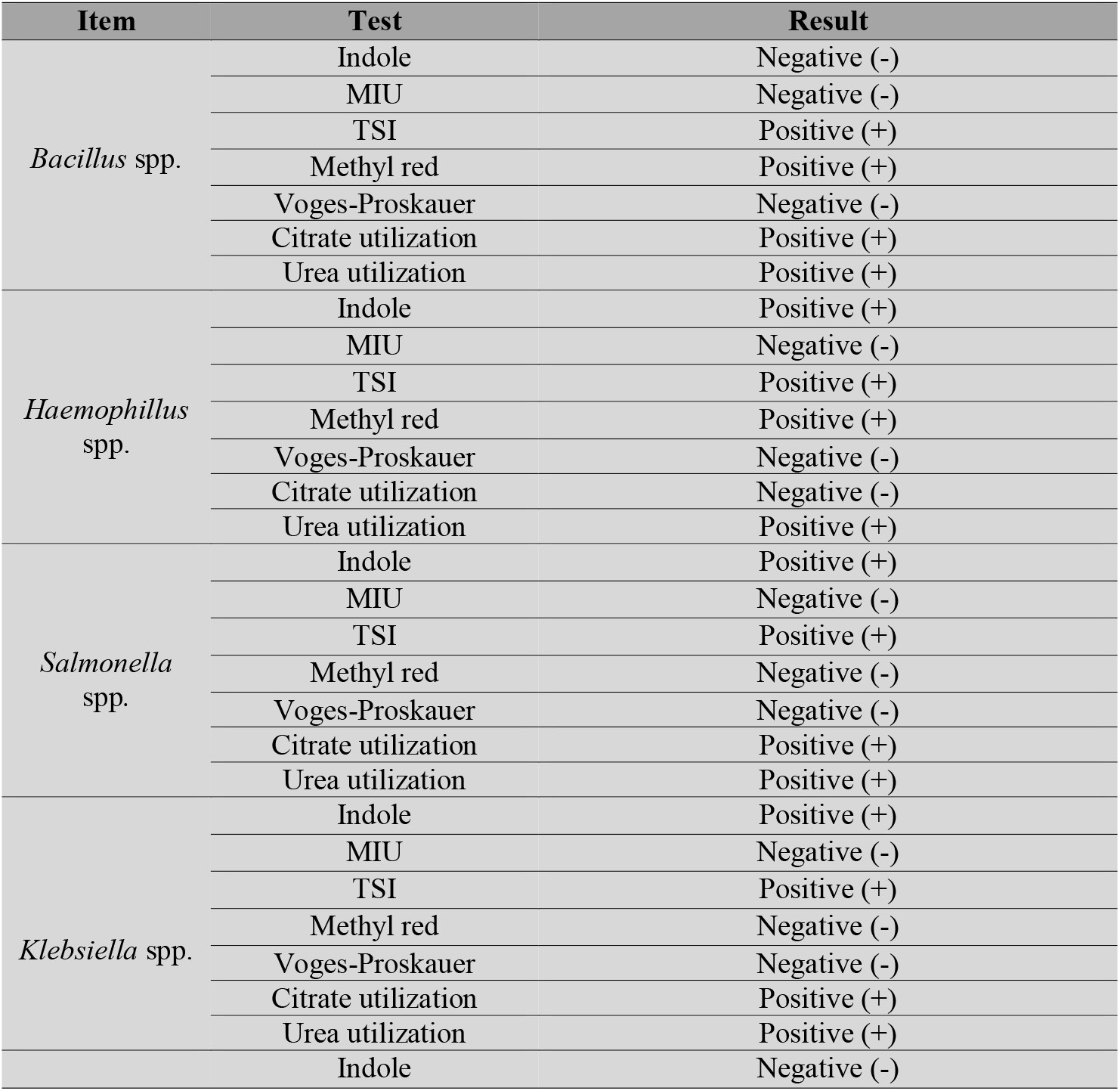
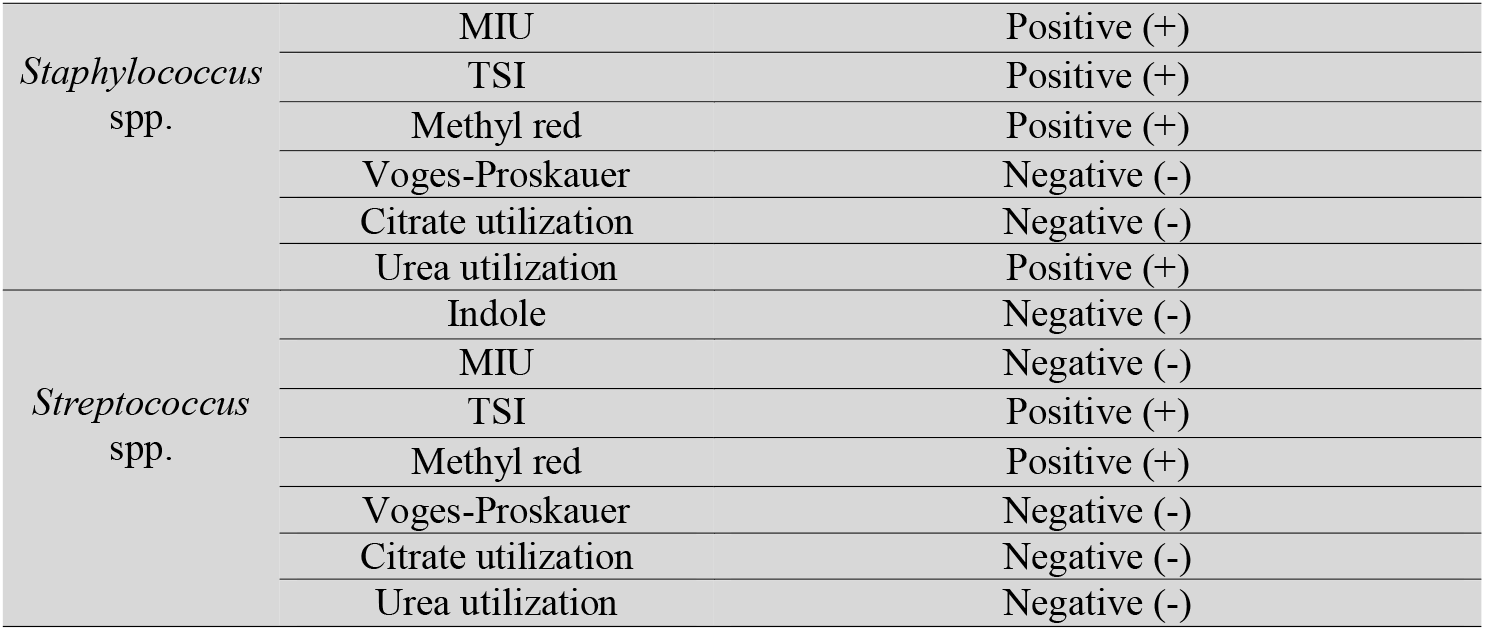
Biochemical properties of isolated bacterial species.

#### Results of antibiotic susceptibility test

A total of six isolates (Bacillus spp., Haemophillus spp., Salmonella spp., Klebsiella spp., Staphylococcus spp., Streptococcus spp.) were subjected to antibiotic sensitivity assay. Antibiotic sensitivity test showed that all the isolated bacterial species were sensitive to ciprofloxacin. On the other hand, all the isolates were resistant to cefixime with the exception of Haemophillus spp. Bacillus spp., Haemophillus spp., Salmonella spp., and Streptococcus spp. were all resistant to the antibiotic azithromycin. While Staphylococcus spp. was sensitive to the antibiotic azithromycin, Klebsiella spp. was intermediate to it. Gentamycin sensitivity ranged among all species of isolated bacteria, while Staphylococcus spp. only showed intermediate sensitivity. As a result, three bacterial species—Salmonella, Bacillus, and Streptococcus— showed 50% resistance to these popular antibiotics, while the other three—Staphylococcus, Klebsiella, and Haemophilus—showed 30% resistance. The results of the antibiotic sensitivity assay are presented in **Table 4**.

**Table 4.**
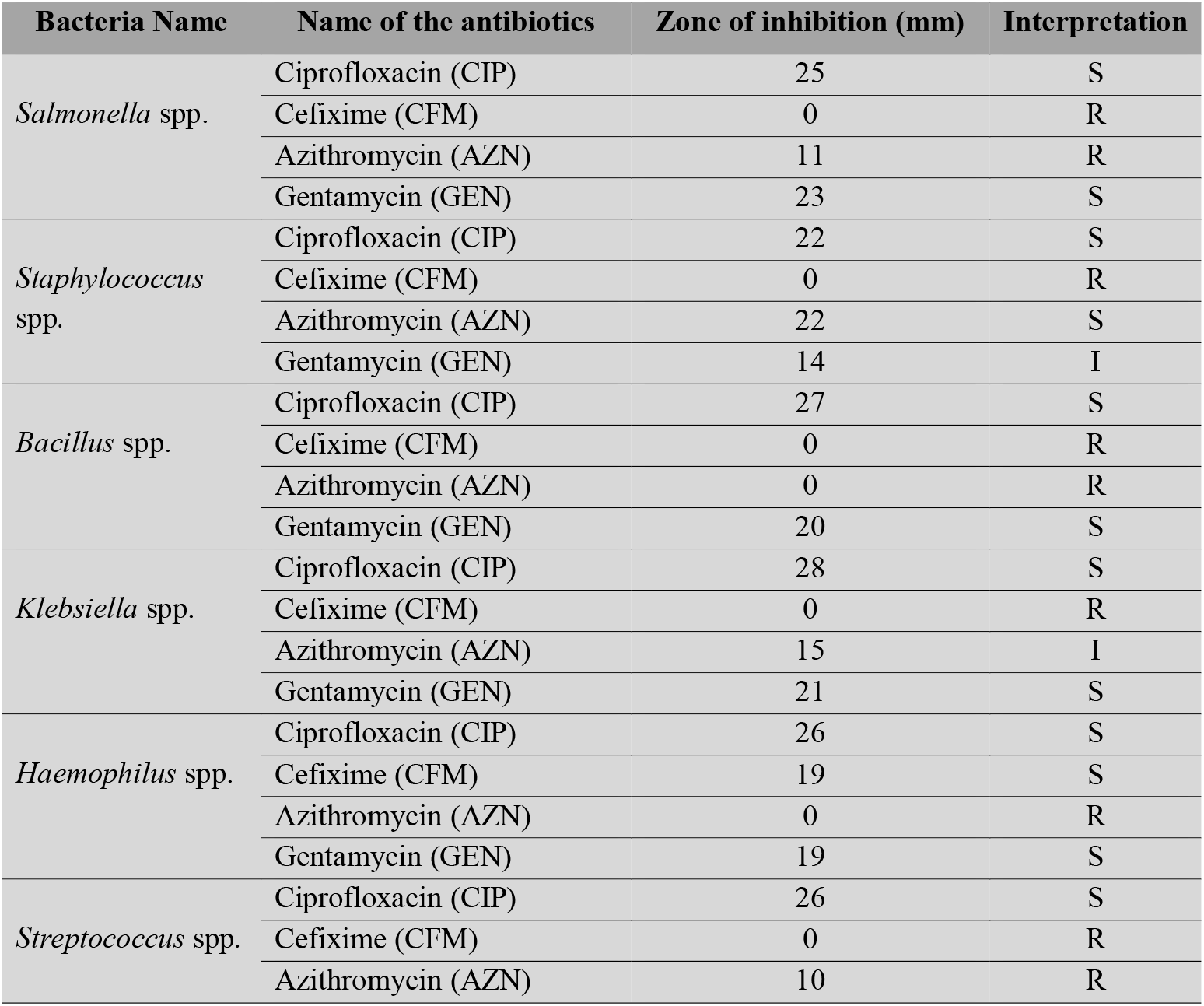
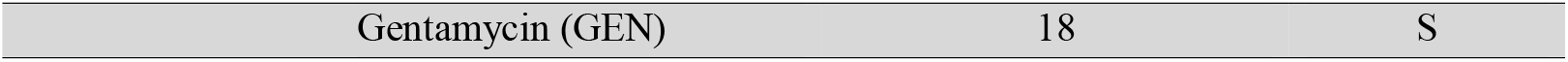
Antibiotic sensitivity test against isolated bacteria.

## Discussion

In the past ten years, there has been a rise in awareness of the role of street food that plays in a number of issues related to public health and hygiene [18]. Perhaps the area that has attracted the most attention and gained the greatest understanding is how these street foods contribute to the spread and acquisition of food-borne diseases. There has been a worldwide rise in the prevalence of illnesses that are transmitted by food [19]. It is a widely held belief at this point that a significant number of incidents of food-borne disease are brought about as a direct result of street food sellers improperly preparing and managing the food they sell [19]. The data collected from all across the world on illnesses caused by Salmonella species and Campylobacter species [20] provide some of the most convincing evidence to date. Both species are responsible for a wide range of diseases, including those related to the gastrointestinal tract and fever [21]. Hemorrhagic colitis, listeriosis, campylobacteriosis, shigellosis, and toxoplasmosis are all caused by germs that are regularly found in food bought off the street [17]. Numerous research studies on the cuisine sold on the streets have previously been carried out in particular regions of Bangladesh [22]. Street foods such Chotpoti, Chanachur, Amra (Spondias mombin), and Jolpai (Elaeocarpus serratus) were contaminated with E. coli, S. aureus, K. pneumoniae, S. typhimurium, S. salmonella, and S. shigella [23]. Staphylococcus spp. and Escherichia coli have been shown to be superior than other forms of bacteria [24].

In our research, we have isolated and characterized food-borne pathogens. We took three samples of Hog plum, Fusca, and Chotpoti as these items are very popular as street food items in Bangladesh. We have isolated six types of bacterial species (Bacillus spp., Haemophillus spp., Salmonella spp., Klebsiella spp., Staphylococcus spp., Streptococcus spp.), where Bacillus spp., and Salmonella spp were available in Chotpoti; Staphylococcus spp. was available in Hog plum; and Haemophillus spp., Klebsiella spp., Staphylococcus spp. and Streptococcus spp. were available in Fusca. We observed that there was not any supremacy of gram-positive or gram-negative bacteria where three were gram-positive and three gram-negative bacterial species.

These bacteria are responsible for various types of disease. Staphylococcus spp. produces a toxin that causes vomiting shortly after being ingested [5]. Staphylococcus bacteria, the microorganisms that cause staph infections, can be found on the skin or in the nose of even healthy people. Deadly staph infections occur when the germs spread to vital organs including the heart, lungs, or circulation [17]. Staph infections are becoming increasingly common in apparently healthy persons, and some of these infections can be fatal [25]. Many diseases are caused by streptococci, which are gram-positive aerobic organisms and causes pharyngitis, pneumonia, wound and skin infections, septicemia, and endocarditis. The affected organ dictates the severity of the symptoms. Rheumatic fever and glomerulonephritis are two possible complications of an infection caused by group A beta-hemolytic streptococci [26]. Klebsiella is a gram-negative bacterium that has been linked to several different forms of healthcare-associated illnesses, such as pneumonia, bloodstream infections, wound or surgical site infections, and meningitis. The site’s primary emphasis is salmonellosis, an infection caused by most strains of Salmonella. Typhoid and paratyphoid fever are both caused by different strains of Salmonella [27]. Although pneumonia is the most common sickness caused by H. influenzae, the bacterium is also capable of causing serious conditions such meningitis and bloodstream infections [28]. Haemophilus influenzae is a bacterium that can result in a wide variety of illnesses [29]. The severity of these infections varies greatly, from the common ear infection to life-threatening ones like those that invade the bloodstream. Though anthrax is still the most well-known Bacillus disease, other Bacillus species have been increasingly linked to a wide variety of infections in recent years [25]. These include abscesses, bacteremia/septicemia, wound and burn infections, ear infections, endocarditis, meningitis, ophthalmitis, osteomyelitis, peritonitis, respiratory and urinary tract infections.

We also performed antibiotic susceptibility tests for these bacterial species with several antibiotics such as ciprofloxacin (CIP), cefixime (CFM), azithromycin (AZN), and gentamycin (GEN). Ciprofloxacin sensitivity was found in all isolated species. Except for Haemophilus spp., all of the bacteria tested positive for cefixime resistance. Bacillus spp., Haemophillus spp., Salmonella spp., and Streptococcus spp. were resistant to azithromycin, whereas Klebsiella spp. were intermediate, and Staphylococcus spp. were susceptible. Gentamycin sensitivity was found in every species, except for Staphylococcus spp., which was merely intermediate. As our experiment revealed, we must exercise caution when consuming these popular street foods. Though these street food items are very delicious and available in our country, we have to prohibit them as soon as possible. To put it another way, vendors selling food on the street should adopt better hygienic practices in how they prepare and serve their products. Otherwise, we will have to suffer very soon, along with our future generations. Besides, there is also a need for more research on this concern in our country for people’s health concerns.

## Conclusion

Street food has been achieved the highest popularity among the young generation right now. Improper personal hygiene can promote the transmission of pathogenic bacteria found in the environment and on people’s hands through these foods. The study’s findings show that, despite their low prices, roadside meals are not healthful owing to unsanitary conditions and unclean vendors. Therefore, these foods sold on the road should be produced according to good hygiene practices and storage procedures should be developed to minimize microbial contamination of the food. Strict public health rules must be established to control the situation. The storage of these fast products should also be closely monitored. The government should take the necessary steps to provide continuing education and awareness on food control and personal hygiene to fast food vendors and consumers.

## Author contributions

MIH and MSA designed the experiment and LBM, MIH, MASMT, MOF, MS and MSA performed, analyzed and interpreted the data, and prepared the manuscript. SRA, KRI, MS and MSA reviewed the manuscript.

## Data availability

All data generated or analysed during this study are included in this article.

## Acknowledgments

The authors would like to thank the Department of Biotechnology and Genetic Engineering, Bangabandhu Sheikh Mujibur Rahman Science and Technology University, Gopalganj, Bangladesh, for providing the research facilities.

